# Comparing Lipinski’s Rule of 5 and Machine Learning Based Prediction of Fraction Absorbed for Assessing Oral Absorption in Humans

**DOI:** 10.1101/2024.08.20.608791

**Authors:** Urban Fagerholm, Sven Hellberg, Jonathan Alvarsson, Morgan Ekmefjord, Ola Spjuth

**Affiliations:** Prosilico AB, Lännavägen 7, SE-141 45 Huddinge, Sweden; Department of Pharmaceutical Biosciences and Science for Life Laboratory, Uppsala University, Box 591, SE-751 24 Uppsala, Sweden

## Abstract

**Background:** The influential Lipinski’s Rule of 5 (Ro5) describes molecular properties important for oral absorption in humans. According to Ro5, poor absorption is more likely when 2 or more of its criteria (molecular weight (MW) >500 g/mol, calculated octanol-water partition coefficient (clog P) >5, >5 hydrogen bond donors (HBD) and >10 hydrogen bond acceptors (HBA)) are violated. Earlier evaluations have shown that many drugs are sufficiently well absorbed into the systemic circulation despite many Ro5-violations. No evaluation of Ro5 *vs* fraction absorbed (f_a_) has, however, been done.

**Methods:** Datasets of orally administered drugs violating and not violating Ro5 and with available human clinical f_a_-values were assembled, and contrasted to machine learning based predictions using the ANDROMEDA prediction software having a major MW-domain of 150-750 g/mol.

**Results:** 129 Ro5-violent compounds (29 with MW>1000 g/mol) were found, 59 of which had f_a_-values (42 % mean f_a_). 34 % and 66 % of compounds were predicted as having f_a_≤10% and >10-30 % respectively, which was in good agreement with measured f_a_ of 37 % and 63 %. The f_a_ for all compounds with f_a_≤5 and ≤10 % were correctly predicted. For compounds with f_a_>30 %, 81 % were predicted to have a f_a_>30 %, but none were predicted to have a f_a_<10 %. The Q^2^ for predicted *vs* observed f_a_ was 0.64. For a set of 77 compounds without Ro5 violation (80 % mean f_a_), all compounds were correctly predicted to have a f_a_ below or above 30 % (Q^2^=0.56). Among these are compounds with poor uptake (<1 % to 7 %).

**Conclusion:** We show that machine learning based predictions of f_a_ are superior to Ro5 for assessing oral absorption obstacles in humans. Too strict reliance on Ro5 may hence constitute a risk. ANDROMEDA predicts f_a_ well, easily and quickly, and also differentiates well between poor and adequate oral uptake for compounds violating and not-violating Ro5. This makes it a valid and useful tool capable of predicting oral absorption in humans with good accuracy and replacing Ro5 for oral absorption assessments.

## Introduction

Classification systems and rules of thumb are common in the field of pharmacokinetic/ biopharmaceutic predictions. Advantages include that they are easy to understand and apply, which increases the possibility of having an impact and creating consensus. Advantages with more complex systems include the possibility to gain mechanistic knowledge and produce numerical, more meaningful, values. Disadvantages with the latter are that they require in-depth knowledge and more data/information and (likely) have lower chance of being accepted as a golden standard. They could also be overly complex and difficult to understand and use and require lots of data. For each data input from a laboratory the uncertainty/variability will increase. For the simpler systems there is a risk of classification errors and oversimplification, and for these it is required that the limits are adequate and well defined/described. Both simple and complex systems are sensitive to variability and uncertainty of data used and *in vitro-in vivo* discrepancies (or species differences) and can/will lose applicability if some required data are unavailable and not possible to produce. The difference between calculated and experimentally determined log P (5.4 and 2.4) of posaconazole exemplifies the uncertainty/variability [1]. For both it is also essential that they are extensively and successfully validated.

The influential Lipinski’s Rule of 5 (Ro5), established molecular properties in 1997, describes molecular properties important for oral absorption (and consequently, oral activity) in humans and states that an orally active drug has no more than one violation of its 4 criteria – molecular weight (MW) >500 g/mol, calculated octanol-water partition coefficient (clog P) >5, >5 hydrogen bond donors (HBD) and >10 hydrogen bond acceptors (HBA) [2]. In other words, poor absorption is more likely when 2 or more of these criteria are violated. There is, however, no definition of poor (e.g. <5%, <10-20 % or <50 % uptake?) and level of likelihood (e.g. >50 % or >95 % likelihood?).

In a review of Ro5 from 2014 by Doak et al. it was highlighted that almost 200 approved oral drugs had a MW above 500 g/mol and that the number of new drugs exceeding that limit increases over time [1]. They also showed that 57, 9 and 54 % of compounds with MW>500 g/mol had poor and good Caco-2 permeability and poor oral bioavailability (F) (<30 %), respectively. Otherwise, they found no correlation between Caco-2 permeability and F. These results demonstrate that the 500 g/mol MW-cutoff is not clearly associated with poor absorption.

*Note:* Log D is also a commonly used parameter in drug discovery. In an analysis by our group we found no correlations between log D and parameters such as F, clearance (lin and log) and half-life (lin and log). Thus, no optimal log D (such as, for example, log D between 0 and 2) for human pharmacokinetics seems to exist.

Hartung et al. evaluated Ro5 violations for 117 oral drugs approved 2018-2022 and found that 19, 17 and 1 % had 3, 2 and 1 violations, respectively [3]. 92, 19, 12 and 42 % had violations of MW, cLog P, HBD and HBA, respectively.

Sufficiently high fraction absorbed (f_a_) and F have been reached for approved oral drugs with MW, clog P and HBD exceeding 1200 g/mol (cyclosporin with 40 % f_a_), 8 (venetoclax with >65 % f_a_) and 8 (k-strophanthoside with 16 % f_a_), respectively [1]. Based on these findings it is clear that too strict reliance on Ro5 may cause lost opportunities [2].

To our knowledge, an analysis of Ro5 related f_a_ has not been done (studies have focused on Ro5 *vs* F and *in vitro* permeability). Such an analysis/validation would give more insight of the applicability and limitations of Ro5.

Our prediction software ANDROMEDA by Prosilico predicts human clinical pharmacokinetics and biopharmaceutics, including f_a_ (considers permeability, dissolution and efflux by MDR-1, BCRP and MRP2) and F. ANDROMEDA, with a major domain for molecules with MW of 150-750 g/mole, is based on conformal prediction (CP), which is a methodology that sits on top of machine learning methods and produce valid levels of confidence [4], unique data and algorithms and a new human physiologically-based pharmacokinetic (PBPK) model [5]. For a more extensive introduction to CP we refer to Alvarsson et al. 2014 and Arvidsson McShane et al. 2024 [6,7]. ANDROMEDA does not predict from the molecular properties MW, log P, HBD and HBA. It has been validated in several (>20) studies and shown to outperform laboratory methods in accuracy and breadth for compounds within and slightly outside of its major MW-domain [8-13]. Advantages with ANDROMEDA include ability to predict batches with up to 1000 compounds at a time and mix *in silico* and laboratory data.

The main objective of the study was to collect orally administered compounds/drugs violating and not violating Ro5 and with available human clinical f_a_-values, for further evaluation of Ro5 and for evaluation of the absolute and relative (*vs* the simple *in silico* rule Ro5) applicability of ANDROMEDA (an *in silico* system without criteria for classifications and limits) for these compounds.

## Methods

### Compound selection

#### Compounds violating Rule of 5

Fifty-nine found and selected compounds with beyond Ro5-properties and human *in vivo* f_a_-data are shown in Table 1. Maximum MW, log P, HBD and HBA are 1880 g/mol (teicoplanin A2 1), 8.2 (venetoclax), 24 (teicoplanin A2 1) and 33 (vancomycin), respectively. Thirty-four and 2 (fidaxomicin and tenapanor) compounds have 3 and 4 Ro5-violations, respectively.

**Table 1.**
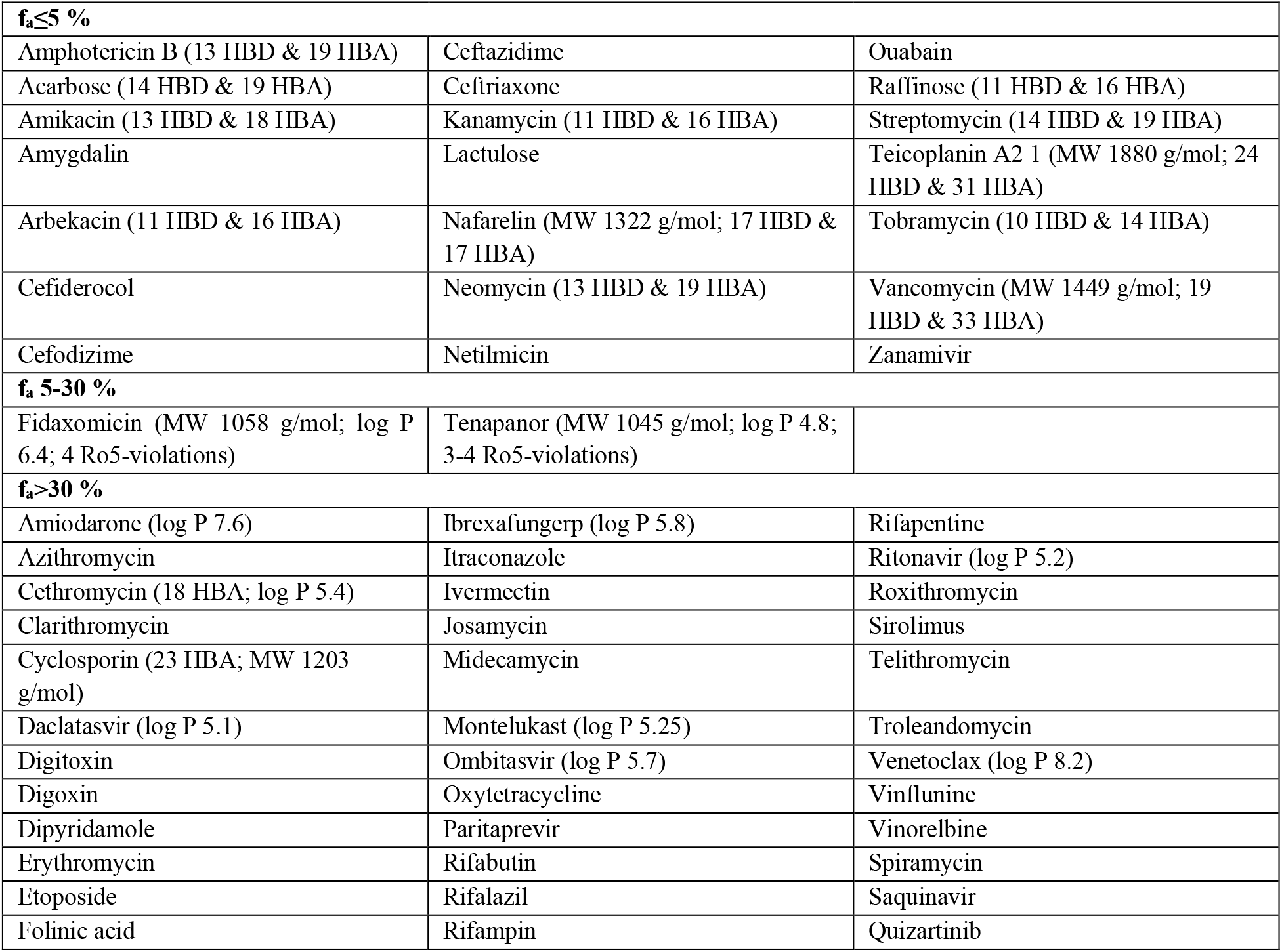
Selected compounds with beyond Ro5-properties and human *in vivo* f_a_-data. Cases with particularly high MW, log P, HBD and HBA are noted.

Prodrugs, hydrolysed compounds and quaternary amines were excluded. Actively absorbed compounds, such as folinic acid and macrolide antibiotics erythromycin, clarithromycin, roxithromycin, azithromycin and telithromycin, were included.

F_a_- and F-values were taken from Pham-Thee et al. [14], Newby et al. [15], Varma et al. [16], FDA approval documents [17], Doak et al. [1], Wikipedia [18], Drugbank [19] and PROSILICOs proprietary data bank. In cases where f_a_ was missing f_a_ was set to >=F.

Compounds were allocated to 3 groups: f_a_≤5 % (n=21), f_a_=5-30 % (n=2) and f_a_>30 % (n=35).

Seventy beyond Ro5-compounds with estimated poor (but not specified) uptake (n=7), >15 % absorption (n=2) and unknown f_a_ (n=61) were excluded from the full analysis, but were part of the evaluation. These include anacetrapib with a log P of 9.3, semaglutide with a MW of 4312 g/mol and 57 HBD and 63 HBA, and imetelstat with a MW of 4610 g/mol and 45 HBD and 107 HBA.

#### Compounds not violating Rule of 5

Seventy-seven compounds with zero (n=70) or 1 (n=7) violations of Ro5, and not included in training sets for absorption in ANDROMEDA, were found and selected. These did not include prodrugs, hydrolysed drugs, quaternary amines, actively absorbed antibiotics, compounds with MW<150 g/mol or >750 g/mol and compounds with uncertain clinical f_a_-values.

F_a_-values were taken from Pham-Thee et al. [14], Newby et al. [15], Varma et al. [16], FDA approval documents [17], Doak et al. [1], Wikipedia [18], Drugbank [19] and PROSILICOs proprietary data bank.

### ANDROMEDA predictions

ANDROMEDA software by Prosilico (version 2.0) was applied to predict f_a_. The dissolution component of f_a_ was based on a default oral dose of 50 mg, and in cases of very low (<few mg) and high (>500 mg) oral doses a correction was made.

Selected compounds in the present study were generally not included in the training sets for the used software version. In the few cases (<6 % of all data for Ro5-violating compounds, but no case for non-violating compounds) they were covered in the underlying models for f_a_, prediction results obtained in previous cross-validation studies were used.

For actively absorbed compounds folinic acid, erythromycin, clarithromycin, roxithromycin, azithromycin and telithromycin uptake was corrected for the active transport contribution. Dose-dependencies for active transport was not considered in the predictions.

## Results & Discussion

### Rule of 5 violating compounds with available fraction absorbed *in vivo* in man

The average f_a_ for compounds violating Ro5 was 42 % (34 % according to predictions). Most (18 of 21) of the compounds with ≤5 % f_a_ had f_a_ of 2 % or less. All of them were successfully predicted to have a f_a_ of 0-2 %. The f_a_ of fidaxomicin (8 %) was also correctly predicted (8 %). These results show that ANDROMEDA accurately predicts the oral uptake of compounds with such properties. For the only compound with f_a_ between 10 and 30 %, tenapanor (f_a_=22 %), ANDROMEDA predicted 46 % uptake. For 36 compounds with >30 % f_a_, ANDROMEDA correctly predicted >30 % f_a_ for 29 (81 %) of them. For the remainder, predicted f_a_ were 10 to 28 %. Thus, ANDROMEDA also predicted adequate f_a_ (ca >10-30 %) well. This makes it very useful for predicting and evaluating whether or not drug candidates are likely to be sufficiently well absorbed following oral dosing in man.

The mean absolute prediction error was 14 %. The Q^2^ (correlation between forward looking predictions and measurements/observations) for predicted *vs* observed f_a_ was ca 0.65 (0.64 when setting f_a_=F when f_a_ was missing; 0.67 when setting f_a_=(100 % + F)/2 in cases where f_a_ was missing) (Figure 1; for 15 compounds with moderate to high degree of absorption f_a_ was set equal to F). Exclusion of folinic acid and mycins with active uptake also resulted in a Q^2^ of 0.6. A Q^2^ of this level was reached despite the complexity and variability of structures, compounds with considerable limitations in permeability and/or dissolution potential, and impact of saturation (active transport and dissolution), intramolecular hydrogen bonds (e.g for macrocycles) and molecular flexibility (e.g. for larger non-cyclic compounds).

**Figure 1.**
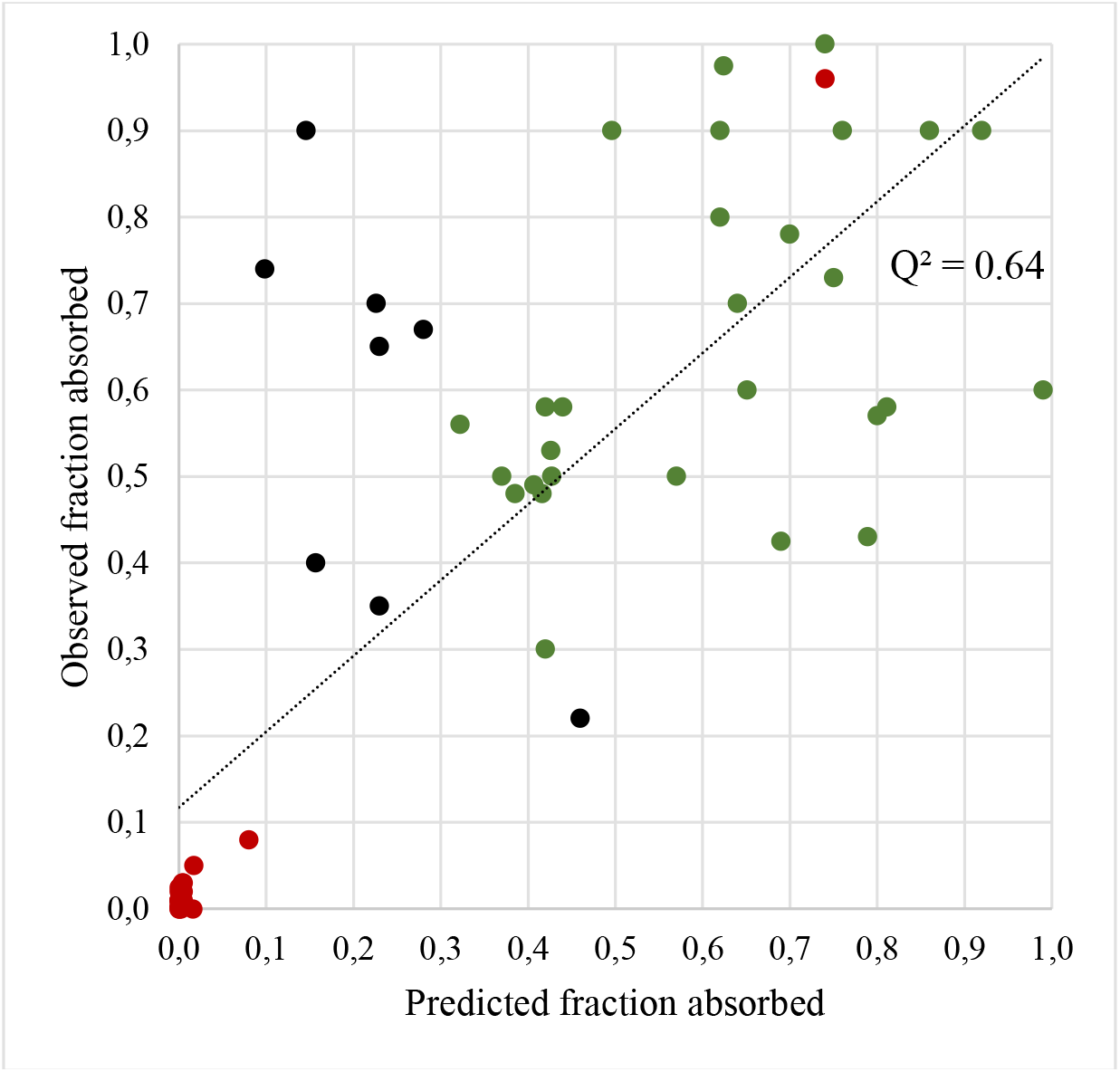
The relationship between predicted and observed *in vivo* f_a_ for beyond Ro5-compounds in humans (n=59). Colors denote compounds with correctly predicted f_a_≤5-10 % (red; n=22) and f_a_>30 % (green; n=29), and incorrect prediction of f_a_ <30< % (black; saturation of efflux was not considered when predicting these; n=8), respectively.

Nine (15 %) of the Ro5-violating compounds belong to *in vivo* BCS I (≥90 % f_a_).

F_a_-values for cyclosporin, dalacatasvir, rifapentine, spiramycin and venetoclax were underpredicted. They are all given at moderate (60 mg) to very high (1000-2000 mg) oral doses and are (or are predicted to be) effluxed in the human intestine. It is possible that saturation of efflux (which can be considered, but was not taken into account in the predictions) might have contributed to these underpredictions.

The f_a_ for tenapanor was overpredicted. The oral dose of tenapanor (50 mg) is lower than for the other compounds with underpredicted f_a_.

Digoxin and digitoxin (with almost complete estimated oral uptake) were the two compounds that deviated most (underpredictions). In Caco-2 experiments, digoxin has moderate permeability and its f_a_ has also been underpredicted (ca 50 % f_a_) using this assay [1,20]. The underpredictions do not seem to be explained by saturation of solubility and/or efflux since they are given at oral doses < 1 mg.

It should be noted that the predictive accuracy also depends on how well *in vivo* f_a_ has been derived.

### Rule of 5 violating compounds without fraction absorbed values in man

The predicted f_a_ for tacrolimus (estimated f_a_>15 %) and velpatasvir (estimated f_a_>25-30 %) were 38 and 29 %, respectively.

The predicted f_a_ for 7 compounds proposed to have poor oral uptake were 0-49 % (average 19 %).

Sixty-one compounds with unknown f_a_ include colistin (predicted f_a_=0 %), semaglutide (predicted f_a_=0 %; MW=4312 g/mol; observed F =0.8 %), dalbavancin (predicted f_a_=0 %; given intravenously only), desmopressin (predicted f_a_=0 %; observed F=0.08-0.16 %), faldaprevir (predicted f_a_=29 %; good *in vitro* permeability; observed F in animals=26-39 %), cobicistat (predicted f_a_=68 %; observed F in animals=11-33 %), lucifer yellow CH (predicted f_a_=61 %; moderate *in vitro* permeability), mezlocillin (predicted f_a_=1.4 %; proposed zero f_a_ *in vivo;* given intravenously only), micafungin (predicted f_a_=1.1 %; F proposed to be similar to F for anidafungin (2-7 %); given intravenously only), pasireotide (predicted f_a_=29 %; observed low *in vitro* permeability), ortataxel (predicted f_a_=42 %; moderate *in vitro* permeability; observed F in animals=25 %) posaconazole (predicted f_a_=81 %; high *in vitro* permeability; observed F in animals=52 %), SCY-635 (predicted f_a_=14 %; %; moderately low *in vitro* permeability; observed F in animals=11-23 %), stevioside (predicted f_a_=0 %; limited permeability), vedroprevir (predicted f_a_=21 %; low *in vitro* permeability; observed F in animals=49-123 %), voclosporin (predicted f_a_=15 %; assumed to be similar to cyclosporin with 40 % f_a_), MK-0616 (predicted f_a_=10 %; observed F =0.2%), terlipressin (predicted f_a_=0 %; given intravenously only), cetrorelix (predicted f_a_=0 %; given subcutaneously only), leuprorelin (predicted f_a_=3 %; given intravenously only), mezlocillin (predicted f_a_=1.4 %; given intravenously only), micafungin (predicted f_a_=1.1 %; given intravenously only), bivalirudin (predicted f_a_=0 %; given intravenously only), satoreotide trizoxetan (predicted f_a_=0 %; given intravenously only), cholecystokinin octapeptide (CCK-8) (predicted f_a_=10 %; observed F =0.2%), moxalactam (predicted f_a_=14 %; given intravenously only), valspodar (predicted f_a_=16 %; given orally), voclosporin (predicted f_a_=15 %; given orally), temsirolimus (MW 1030) (predicted f_a_=12 %; given intravenously only), motixafortide (predicted f_a_= 0 %; given subcutaneously only), rezafungin (predicted f_a_= 8.5%; given intravenously only), zilucoplan (predicted f_a_= 0 %; MW 3562 g/mol; given subcutaneously only), imetelstat (predicted f_a_= 0 %; MW 4610 g/mol; given intravenously only), ceftobiprole medocaril sodium (predicted f_a_= 0.7 %; given intravenously only) and omaveloxolone (predicted f_a_=62 %; observed F=8%?). Overall, predicted f_a_-values for compounds with reference data were in line with available F in animals, *in vitro* permeability and chosen routes of administration (oral, intravenous only or subcutaneous only).

The lowest log P reported for a compound used in this study was -6.6 (for neomycin). The results show that ANDROMEDA is applicable for compounds with large MW (up to at least 4600 g/mol) and very low (<-6) and high log P (>9).

### Predicted molecular signatures

ANDROMEDA produces signatures for each model parameter for each predicted compound. Figure 2 shows the molecular regions contributing to increasing and decreasing the passive permeability (left) and fraction dissolved *in vivo* (right) for teicoplanin A2 1 (MW 1880 g/mol; log P=-2.3; 24 HBD; 31 HBA), which has zero predicted and observed f_a_. The more intense the color and more colored molecular regions, the more information from training set data were useful for predictions. Predicted f_a_ and Figure 2 show that ANDROMEDA is applicable for predicting the biopharmaceutics of complex molecules of this size.

**Figure 2.**
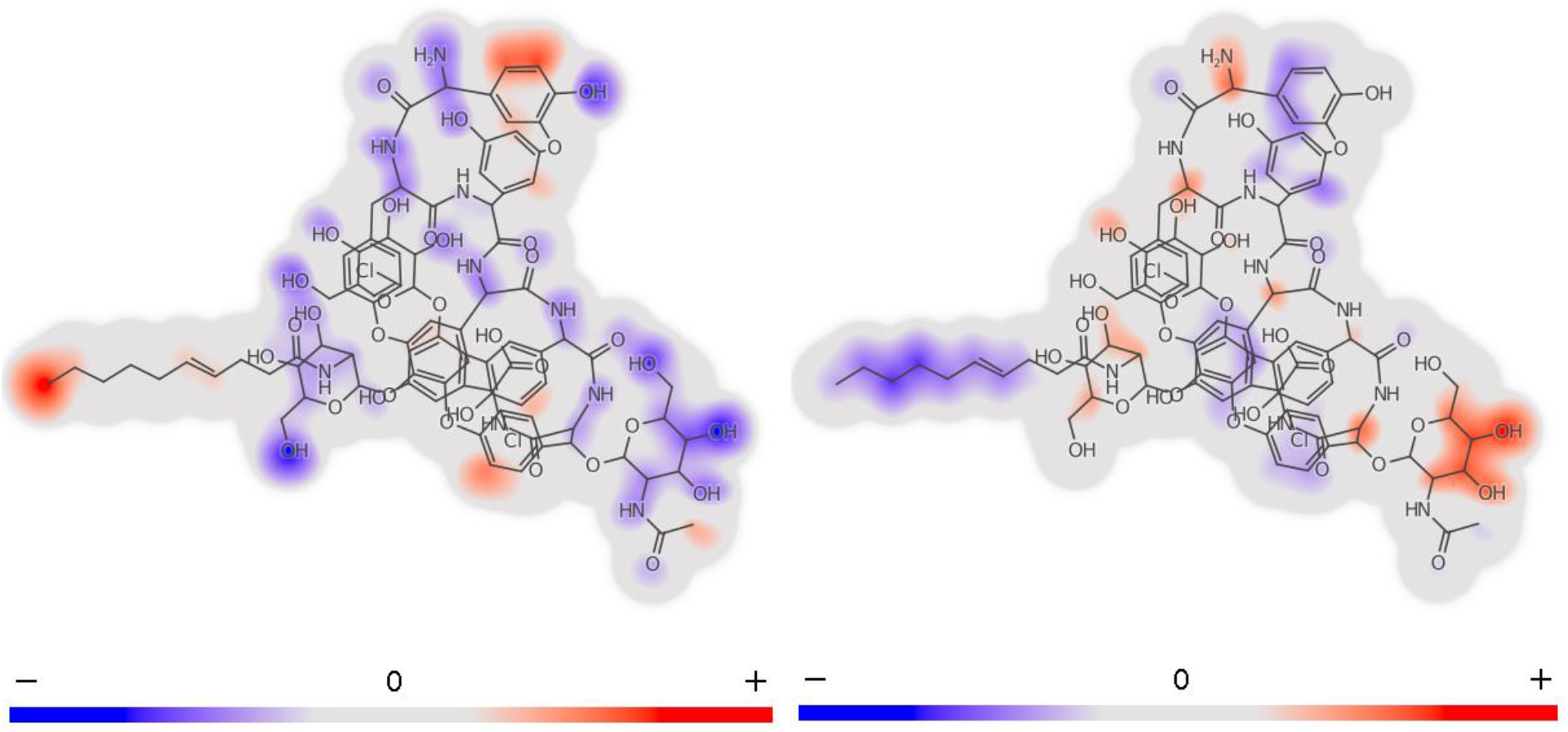
Molecular regions contributing to increasing (red) and decreasing (blue) the passive permeability (left) and fraction dissolved *in vivo* (right) for teicoplanin A2 1.

### ANDROMEDA *vs* Ro5

According to the simple Ro5 all of the 129 selected compounds (29 of them have a MW exceeding 1000 g/mol) should be poorly/insufficiently absorbed following oral administration in humans. No clear definition of poor/insufficient has been done, which is a limitation. In contrast, the more complex ANDROMEDA software predicted that a) 44 (34 %) of these would be insufficiently absorbed (f_a_≤10 %; 39 with f_a_≤5 %), b) 85 (66 %) would be sufficiently absorbed (>10-30 %), c) 34 (26 %) has a f_a_>50 %), and d) 2 (2 %) has a f_a_>90 % (maximum 99 %). The corresponding percentages for observed f_a_ ≤10 and >10-30 % were 37 and 63 % (n=59), respectively, which is in good agreement with ANDROMEDA predictions.

Not only does ANDROMEDA predict f_a_ well (producing f_a_-values and molecular structure information), it also differentiates well between poor (≤5-10 %) and adequate (>10-30 %) oral uptake for beyond Ro5-compounds. It is also devoid of the lab variability/uncertainty factor and is easy to apply and efficient. This makes it a valid and useful tool capable of replacing the Ro5.

### Limitations with *in vitro* methods and data

In an analysis by our group we found no correlations between aqueous solubility (not even for dose-corrected solubility) and *in vivo* dissolution and F. Other limitations with laboratory methods include binding to material and low recovery, *in vitro-in vivo* differences (including active transport), variability (as shown above) and measurement/detection limitations. For example, of the compounds in this study the following have not been able to quantify in an *in vitro* permeability assay (≤LOQ [1,21]): amiodarone, amygdalin, cefodizime, ceftriaxone, dalbavancin, danoprevir, folinic acid, octreotide, ombitasvir, paclitaxel, suramin, terlipressin, vancomycin and vincristine. The following compounds are practically insoluble in water (<LOQ): anidulafungin, azithromycin, bivalirudin, elbasvir, everolimus, posaconazole, quinupristin, ridaforolimus, rifaximin, vedroprevir, velpatasvir, venetoclax and voclosporin. ANDROMEDA is able to predict the f_a_ by permeation and dissolution for all these and all the many BCS IV (low permeability and low solubility according to the Biopharmaceutics Classification System) drugs in this study, which is another advantage.

### Compounds not violating Rule of 5

For 77 compounds with zero or 1 violation of Ro5 the average f_a_ was 80 % (72 % according to predictions) (Figure 3). All these compounds were correctly predicted to have a f_a_ below or above 30 % and the Q^2^ for predicted *vs* observed f_a_ was 0.56 (similar to that for compounds violating Ro5; however, relatively few low f_a_-compounds; a Q^2^ of this level is somewhat lower than that obtained for a larger dataset in previous internal validations) (Figure 4). The mean absolute and relative prediction errors were 14 % (similar to that for compounds violating Ro5) and 1.30-fold, respectively. Dose-dependencies for active transport and dissolution was not considered in the predictions, which might have influenced the results (over- or underpredictions of f_a_ when high oral doses were given).

**Figure 3.**
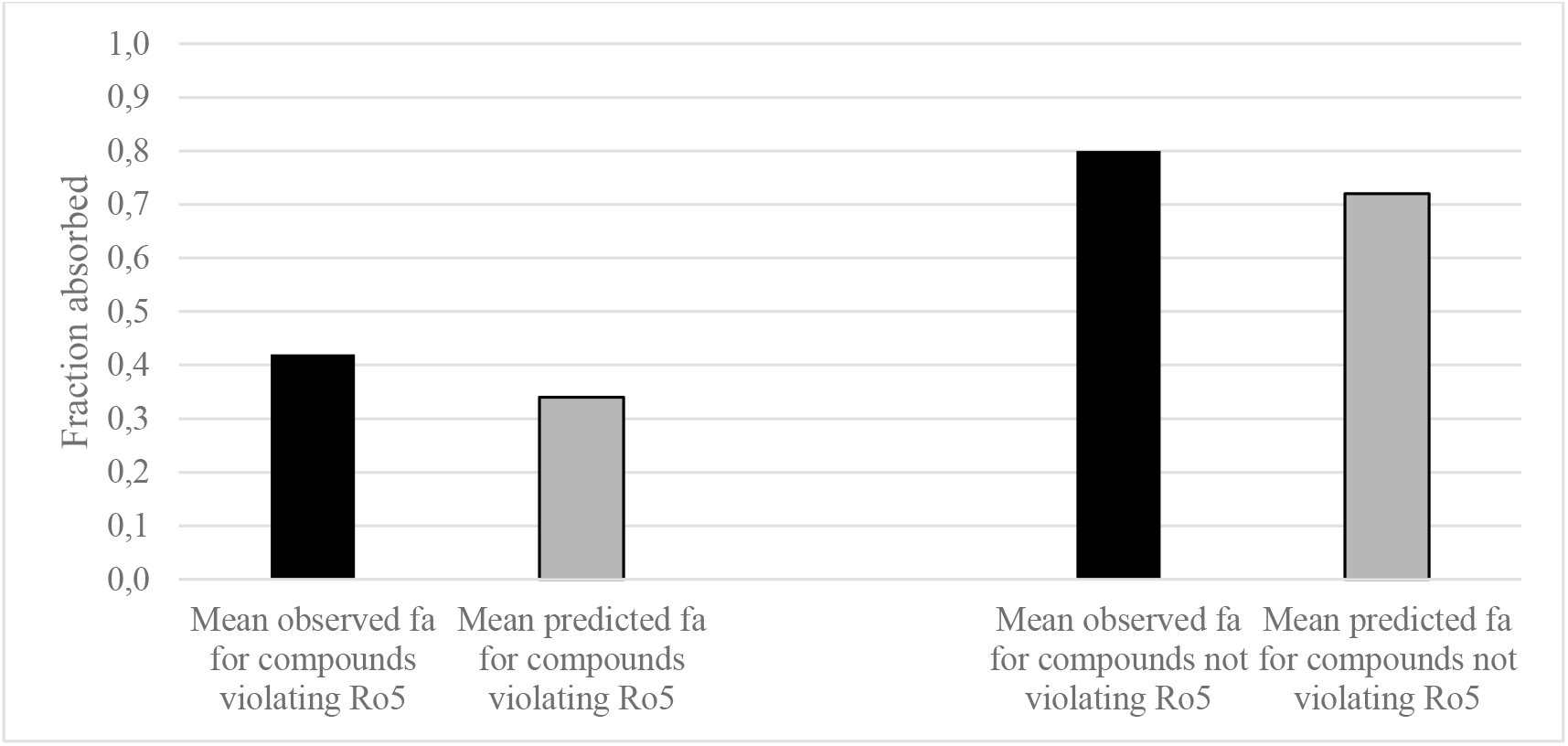
The average observed/estimated and predicted f_a_ for compounds violating or not-violating Ro5.

**Figure 4.**
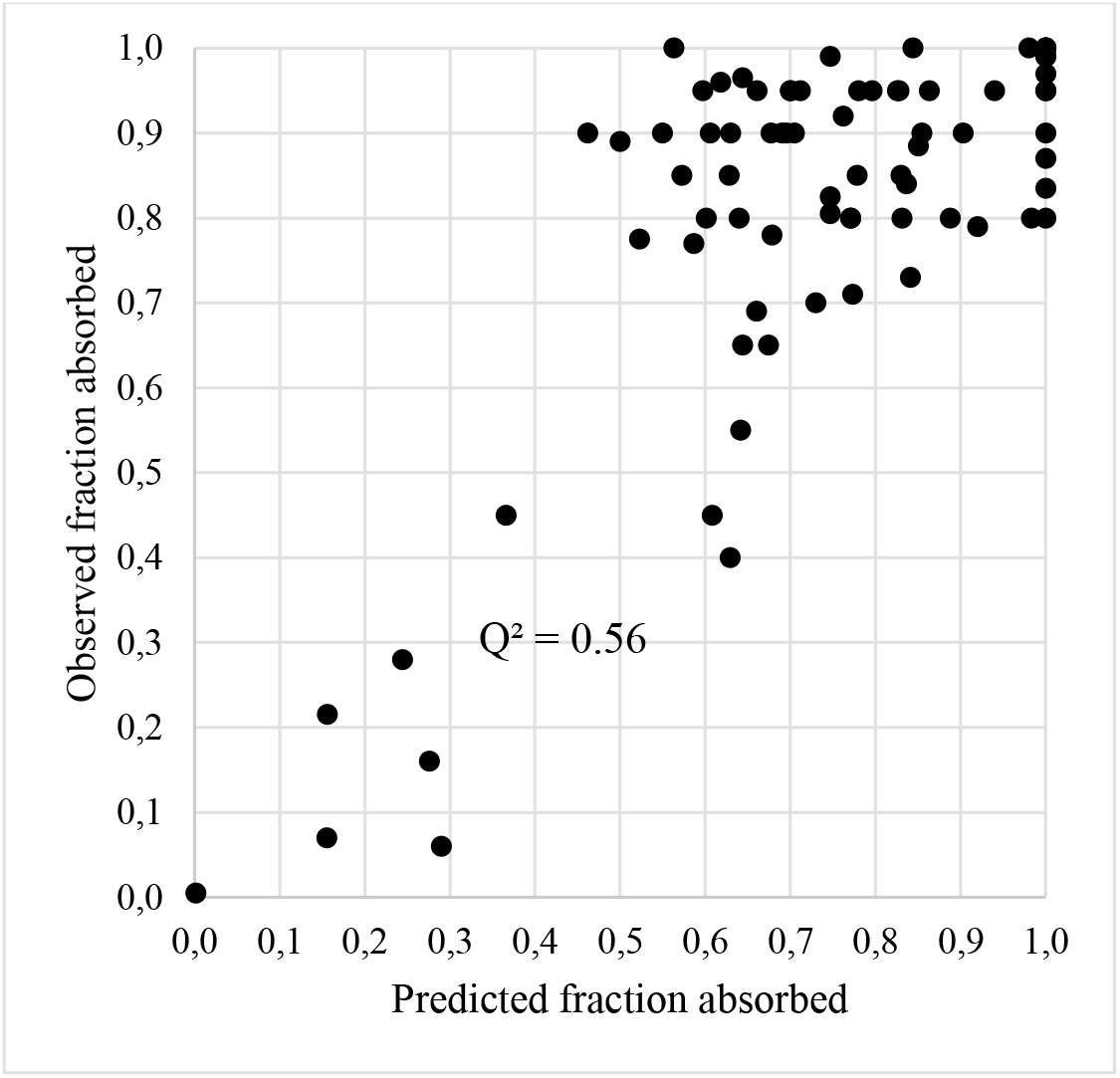
The relationship between predicted and observed *in vivo* f_a_ for 77 compounds not violating Ro5-compounds in humans.

Seven of the compounds had less than 90 % *in vivo* dissolution at an oral dose of 50 mg (*in vivo* BCS III and IV) and 38 (every other compound) belonged to *in vivo* BCS I (≥90 % f_a_).

Three (4 %) and 6 (8 %) of the 77 compounds not violating Ro5 had a f_a_<10 and 30 %, respectively. The lowest f_a_ for compounds with no or one Ro5-violation were <1 and 6 %, and 7 and 16 %, respectively. This shows that poor uptake might occur in humans despite being on the right side of the cut-off of Ro5.

## Conclusion

We show that machine learning based predictions of f_a_ are superior to Ro5 for assessing oral absorption obstacles in humans. Too strict reliance on Ro5 may hence constitute a risk, including lost opportunities and unnecessary failures. ANDROMEDA predicts f_a_ well, easily and quickly, and also differentiates well between poor and adequate oral uptake for compounds violating and not-violating Ro5. This makes it a valid and useful tool capable of predicting oral absorption of small drugs and drug candidates in humans with good accuracy and replacing Ro5 for oral absorption assessments.

## References

1. Doak BC, Over B, Giordanetto F & Kihlberg J. (2014). Review - Oral druggable space beyond the Rule of 5: Insights from drugs and clinical candidates. Chem. & Biol. 21, 1115–1142.

2. Lipinski CA, Lombardo F, Dominy BW & Feeney PJ. (1997) Experimental and computational approaches to estimate solubility and permeability in drug discovery and development settings. Adv. Drug Deliv. Rev. 23, 3–25.

3. Hartung IV, Huck BR & Crespo A. (2023). Rules made to be broken. Nat. Rev. Chem. 7, 3–4.

4. Vovk V, Gammerman A & Shafer G. (2005). Algorithmic learning in a random world. Springer Science & Business Media.

5. Fagerholm U, Hellberg S & Spjuth O. (2021). Advances in predictions of oral bioavailability of candidate drugs in man with new machine learning methodology. Molecules 26, 2572.

6. Alvarsson J, Arvidsson McShane S, Norinder U & Spjuth O. (2021). Predicting with confidence: using conformal prediction in drug discovery. J. Pharm. Sci. 110, 42–49.

7. Arvidsson McShane S, Norinder U, Alvarsson J, Ahlberg E, Carlsson L & Spjuth O. (2024). CPSign: conformal prediction for cheminformatics modeling. J. Cheminf. 16, 75.

8. Fagerholm U, Spjuth O & Hellberg S. (2022). The impact of reference data selection for the prediction accuracy of intrinsic hepatic metabolic clearance. J. Pharm. Sci. 111, 2645–2649.

9. Fagerholm U, Hellberg S, Alvarsson J & Spjuth O. (2021) In silico prediction of volume of distribution of drugs in man using conformal prediction performs on par with animal data-based models. Xenobiot. 51, 1366–1371.

10. Fagerholm U, Hellberg S, Alvarsson J & Spjuth O. (2021). In silico predictions of the human pharmacokinetics/toxicokinetics of 65 chemicals from various classes using conformal prediction methodology. Xenobiot. 51, 1366–1371.

11. Fagerholm U, Hellberg S, Alvarsson J & Spjuth O. (2022) In silico predictions of the gastrointestinal uptake of macrocycles in man using conformal prediction methodology. J Pharm. Sci. 1111, 2614–2619.

12. Fagerholm U, Hellberg S, Alvarsson J & Spjuth O. (2022) Using the ANDROMEDA by Prosilico software for prediction of the human pharmacokinetics of 4 compounds of natural origin -colistin, curucumin, UCN-01 and voclosporin. BioRxiv.

13. Fagerholm U, Hellberg S, Alvarsson J & Spjuth O. (2023) In silico prediction of human clinical pharmacokinetics with ANDROMEDA by Prosilico: Predictions for an established benchmarking data set, a modern small drug data set, and a comparison with laboratory methods. Altern. Lab. Anim. 51, 39–54.

14. Pham-The H, Garrigues T, Bermejo M, González-Álvarez I, Cruz Monteagudo M & Ángel Cabrera-Pérez M. (2013). Provisional classification and in silico study of biopharmaceutical system based on Caco-2 cell permeability and dose number. Mol Pharmaceut. 10, 2445–2461.

15. Newby D, Freitas AA & Ghafourian T. (2015). Decision trees to characterise the roles of permeability and solubility on the prediction of oral absorption. Eur. J. Med. Chem., 90, 751–765.

16. Varma MVS, Obach RS, Rotter C, Miller HR, Chang G, Steyn SJ, El-Kattan A & Troutman MD. (2010). Physicochemical space for optimum oral bioavailability: Contribution of human intestinal absorption and first-pass elimination. J. Med. Chem. 53, 1098–1108.

17. FDA - https://www.fda.gov/drugs/.

18. Wikipedia - https://sv.wikipedia.org/wiki/.

19. Drugbank - https://go.drugbank.com/.

20. Matsson P, Bergström CAS, Nagahara N, Tavelin S, Norinder U & Artursson, P. (2005). Exploring the role of different drug transport routes in permeability screening. J. Med. Chem. 48, 604–613.

21. Keefer CE, Chang G, Di L, et al. (2023). The comparison of machine learning and mechanistic in vitro-in vivo extrapolation models for the prediction of human intrinsic clearance. Mol. Pharm. 6, 5616–5630.

